# Layer-by-Layer Polymeric Films: A Novel Approach to Buccal GLP-1 Delivery

**DOI:** 10.64898/2026.01.19.700335

**Authors:** Eleftheria Pantazoglou, Floriane Bahuon, Amalie Kjær Andresen, Matteo Tollemeto, Zhongyang Zhang, Ioannis Tzitzigiannis, Nazanin Zanjanizadeh Ezazi, Margarida M. A. Sacramento, João F. Mano, Gavrielle R. Untracht, Peter E. Andersen, Marco van de Weert, Ragna Berthelsen, Stephen T. Buckley, Leticia Hosta-Rigau, Jette Jacobsen, Line Hagner Nielsen

## Abstract

Buccal delivery offers a promising alternative to oral drug administration by enabling direct systemic absorption and avoiding first-pass metabolism. Multilayer polymeric films represent a promising strategy for the sequential delivery of drug and absorption enhancer in the oral cavity. Here, dual- and triple-layer films were fabricated via slot-die coating, incorporating a GLP-1 receptor agonist (GLP-1-RA) and the penetration enhancer sodium glycodeoxycholate (GDC). These were co-loaded in dual-layer films or compartmentalized in triple-layer films. Scanning electron microscopy and optical coherence tomography confirmed well-defined, distinct layers with thicknesses suitable for buccal administration (339 ± 10.24 µm and 487 ± 36.5 µm for dual- and triple-layer films, respectively). Both designs exhibited good mucoadhesion and mucosal compatibility, and preserved the secondary structure of GLP-1-RA. *In vitro* release studies showed rapid diffusion of GDC and GLP-1-RA from dual-layer films, whereas triple-layer films enabled sustained, sequential release of GDC and GLP-1-RA. *Ex vivo* porcine buccal mucosa studies showed higher GLP-1-RA and GDC flux from triple-layer films compared to dual-layer films. The films also did not compromise epithelial integrity, in contrast to the direct application of GLP-1-RA and GDC, which caused significant epithelial disruption. These results demonstrate that multilayer film architecture and spatial layering can be harnessed to control release kinetics, maximize peptide penetration, and minimize tissue stress, offering a versatile platform for safe and effective peptide delivery.

**Graphical abstract:** 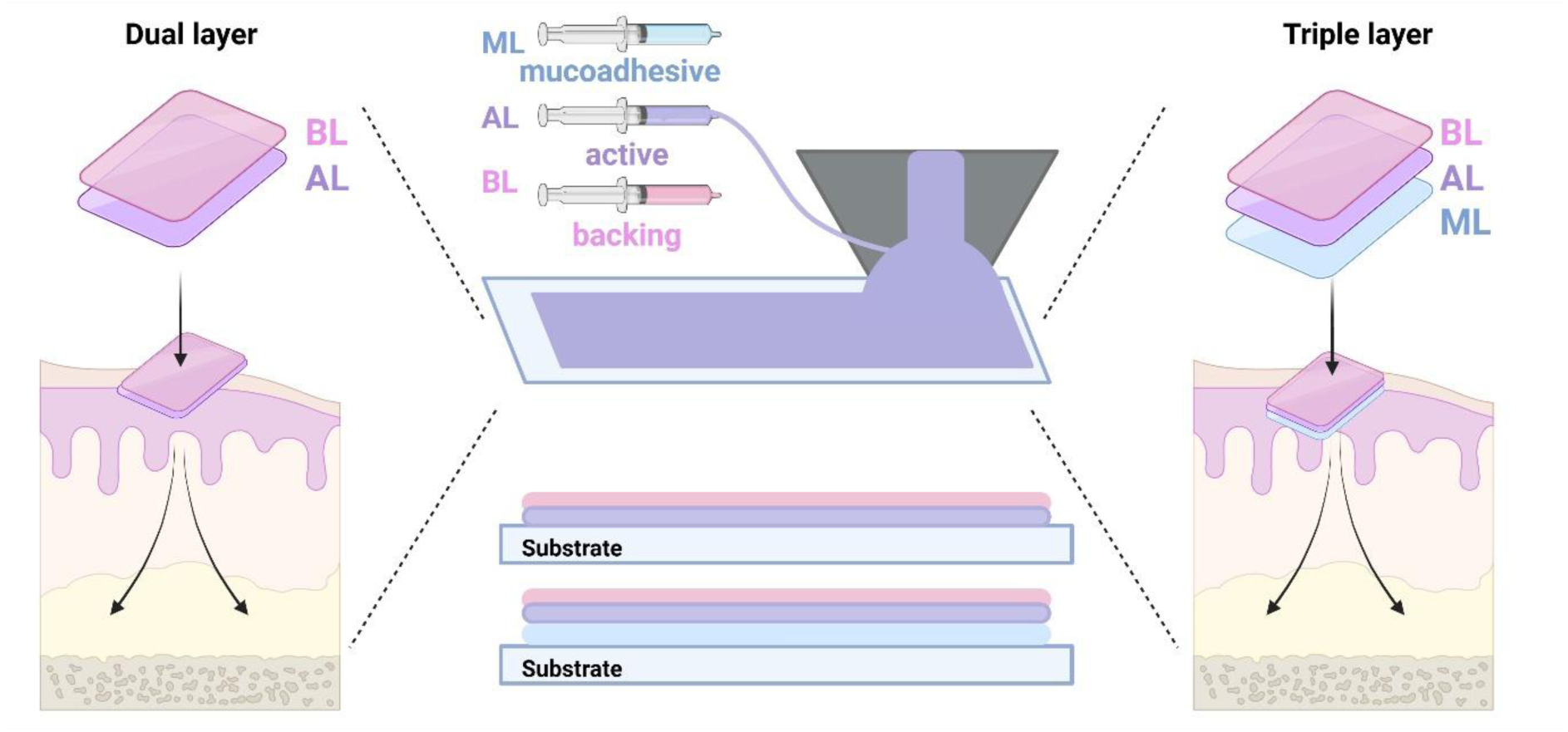

## 1. INTRODUCTION

Peptides such as glucagon-like peptide-1 (GLP-1) are highly effective therapeutic agents due to their potent biological activity and high target specificity.^1,2^ Native GLP-1 is a 30–31 amino acids peptide (approximately 3.3 kDa) with high aqueous solubility but limited stability to enzymatic degradation. Modified analogues, GLP-1 Receptor Agonists (GLP-1 RAs), such as semaglutide (approximately 4.1 kDa) include hydrophobic fatty-acid chains and amino-acid substitutions that increase half-life but reduce solubility, particularly below pH 6.^3–6^ However, despite these advantageous pharmacokinetic modifications, GLP-1-RAs still face significant formulation and delivery hurdles.^7,8^

Their clinical use is limited by rapid enzymatic degradation and poor permeability across biological barriers, especially in oral and mucosal delivery routes.^9,10^ Developing advanced drug delivery systems which protect GLP-1 from degradation and maintain its structural stability is therefore essential to ensure sustained therapeutic activity and improved absorption. Moreover, as GLP-1 is used for chronic conditions such as type 2 diabetes, patient-friendly routes of administration are essential in the design of its drug delivery systems. Currently available oral GLP-1 formulations exhibit very low bioavailability (approximately 1%), highlighting the need for alternative delivery routes.^11^

The buccal route offers a promising alternative, providing direct systemic absorption through the non-keratinized buccal mucosa. It has a near-neutral pH of approximately 6.8, exhibits low enzymatic activity, and is highly vascularized enabling rapid drug uptake.^12,13^ Recent advancements in buccal drug delivery have introduced a range of innovative strategies, including microneedle-based patches and penetration-enhancing formulations.^14^ By applying drug-loaded microneedle patches to the buccal mucosa, insulin and human growth hormone were rapidly delivered.^15^ A needle-free microjet device has also been used to deliver vaccines to the buccal mucosa, achieving robust local and systemic antibody responses in animal models.^16^ Recently, a suction patch inspired by octopus suckers has been developed. This device stretches the buccal mucosa and, with the use of penetration enhancers, significantly improved bioavailability of the peptide desmopressin was achieved.^17^ These findings suggest that buccal administration represents a promising route for GLP-1 delivery; however, current delivery devices are technologically complex, which pose challenges for large-scale manufacturing and clinical translation.

Thin mucoadhesive films are a versatile platform for buccal drug delivery, enabling rapid or controlled release, and good contact with the mucosal surface.^18^ Typically composed of mucoadhesive polymers, these films are designed to adhere to the buccal mucosa and enable unidirectional release of drugs while minimizing drug loss into the oral cavity.^19^ Bilayer films have been investigated for drug delivery applications. For example, mucoadhesive bilayer films made of hydroxypropyl methylcellulose (HPMC) for triamcinolone acetonide demonstrated improved permeability compared to monolayer films, particularly when a backing layer was included to promote unidirectional release.^20^ Furthermore, a multilayered mucoadhesive “nanofiber-on-foam-on-film” system combining chitosan nanofibers, a peptide-loaded cellulose foam, and a saliva-repelling backing film has been developed. This showed improved mucoadhesion, rapid peptide release, and enhanced buccal penetration of desmopressin compared to a commercial tablet.^21^

Despite recent advances, current buccal delivery systems remain limited by payload capacity (often too low to match clinically relevant doses), delivery rate, peptide stability, patient comfort, and scalable fabrication.^22,23^ This highlights the need for a platform capable of efficiently delivering macromolecules in a stable, minimally invasive manner— a challenge addressed in the present study. Building on our earlier proof-of-concept with slot-die-coated pectin monolayer films for drug delivery^24^, we expanded this scalable fabrication approach to dual- and triple-layer prototypes incorporating a GLP-1-RA and a penetration enhancer, to investigate differences in release behavior and permeability through porcine buccal tissue. Importantly, these multilayer films were designed with a unidirectional release architecture, in which all surfaces except the one in contact with the buccal mucosa are protected by the backing layer, thereby minimizing drug loss and ensuring targeted delivery at the tissue interface.

## 2. MATERIALS & METHODS

Glycerol (GLY, ≥99.5%), Pluronic® F-127 (PLU; MW 12,600 Da), poly(acrylic acid) (PAA; MW 250,000 Da, 35 wt.% in H₂O), mucins from bovine submaxillary glands (Type I-S), sodium glycodeoxycholate (GDC), 10% neutral buffered formalin (NBF), Eosin Y 1% alcoholic solution, and glacial acetic acid were all purchased from Sigma-Aldrich (St. Louis, MO, USA). Pertex mounting medium was obtained from Histolab Products AB (Gothenburg, Sweden). Carboxymethyl cellulose sodium salt (NaCMC; average MW 90,000 Da) was obtained from Thermo Fisher Scientific (Waltham, MA, USA). Kollicoat® Protect (KCP; MW 45,000) was purchased from BASF SE (Ludwigshafen, Germany), while hydroxypropyl cellulose (HPC) grade L (MW 140,000) was from Nippon Soda Co., Ltd. (Tokyo, Japan). Glucagon-like peptide-1 receptor agonist (GLP-1 RA) was kindly provided by Novo Nordisk A/S (Måløv, Denmark). Ultrapure water (18.2 MΩ·cm; Milli-Q®, Merck Millipore, Darmstadt, Germany) was used for all experiments. For cell culture experiments, Dulbecco’s Modified Eagle Medium/Nutrient Mixture F-12 (DMEM/F-12), MEM non-essential amino acid solution (100×), fetal bovine serum (FBS), phosphate-buffered saline (PBS; pH 7.4, sterile), penicillin–streptomycin solution (5,000 U/mL), and TrypLE™ Express were obtained from Gibco™ (Thermo Fisher Scientific, Paisley, United Kingdom). L-glutamine and Triton™ X-100 were purchased from Sigma-Aldrich (St. Louis, MO, USA). The CyQUANT™ MTT Cell Viability Assay Kit was supplied by Invitrogen™ (Thermo Fisher Scientific, Waltham, MA, USA). All reagents were used as received without further purification.

### 2.1 Preparation of polymer solutions

Film-forming polymeric solutions were prepared for three distinct layers: the mucoadhesive layer (ML), the active layer (AL), and the backing layer (BL). Triple-layer films consisted of all three layers (ML, AL, BL), whereas dual-layer films were only composed of AL and BL as shown in Figure 1. For each layer, the specific formulation components (as detailed in Table 1) were gradually added to ultrapure water at their designated concentrations. Each component was fully dissolved before adding the next, ensuring a homogeneous, clear solution free of air bubbles and agglomerates. Dissolution was performed on a magnetic stirrer at 350 rpm, with initial heating at 50 °C for 1 h, followed by continued stirring at room temperature until complete solvation.

**Figure 1:**
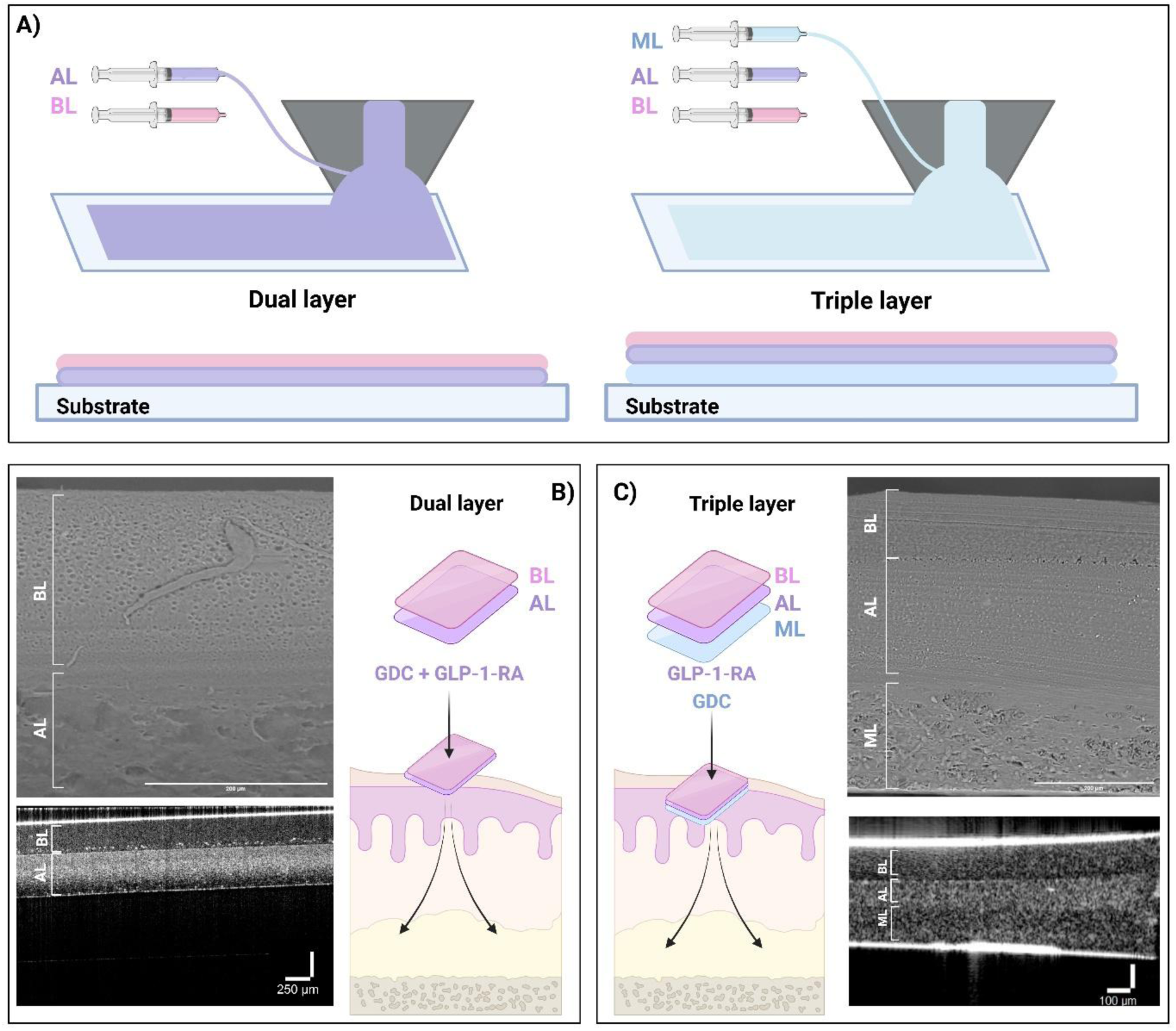
Characterization of dual- and triple-layer films. A) Schematic drawing of the slot-die coating process used to fabricate dual- and triple-layer films. B, C) Schematic representations and corresponding SEM (scale bars represent 200 μm) and OCT images of dual-and triple-layer films (scale bars represent 250 μm (B) and 100 μm (C)).

**Table 1:** Composition of the individual layers (ML: mucoadhesive, AL: active, BL: backing) for the dual- and triple-layer films.

| Film | Layer | Components (mg/mL) |  |  |  |  |  |  |  |
| --- | --- | --- | --- | --- | --- | --- | --- | --- | --- |
|  |  | PLU | HPC | GLY | NaCMC | PAA | KP | GDC | GLP-1-RA |
| 3L | ML | - | - | 50 | 25 | 25 | - | 20 | - |
| 3L | AL | - | - | 50 | 25 | - | 50 | - | 20 |
| 2L | AL | - | - | 50 | 25 | - | 50 | 20 | 20 |
| 2L, 3L | BL | 37.5 | 150 | 20 | - | - | - | - | - |

\* Carboxymethyl cellulose sodium salt (NaCMC), Glycerol (GLY), Pluronic® F-127 (PLU), poly(acrylic acid) (PAA), sodium glycodeoxycholate (GDC), Kollicoat® Protect (KP), hydroxypropyl cellulose (HPC), glucagon-like peptide-1 receptor agonist (GLP-1 RA)

In the dual-layer films, both the GLP-1-RA and the penetration enhancer (GDC) were incorporated into the AL. In contrast, for the triple-layer films, GDC was incorporated into ML while the GLP-1-RA was loaded into AL.

### 2.2 Film fabrication using slot-die coating

All the resulting solutions were cast on the base plate of the film casting apparatus, a slot-die coater (Research Laboratory Coater) from infinityPV ApS (Jyllinge, Denmark) (Fig.1A). For the dual-layer films, the AL solution was cast first and allowed to dry completely. Once dried, the BL solution was cast directly on top of the AL and left to dry. For the triple-layer films, the ML was cast first and dried followed by the casting of the AL. Finally, the BL was cast on top of the AL and dried to complete the triple-layer structure.

For the casting process, a 40 mm slot-die head with a working width of 10 mm and a gap height of 1 mm was used for ML and AL, whereas a 65 mm slot-die head with a working width of 50 mm and a gap height of 1 mm was used for the BL to fully cover the previous layers. The pump speed was 4 mL/min, and the slot-die head speed was 1.2 mm/min. A sheet of polyethylene terephthalate (PET) was used as a liner on the base plate of the slot-die coater. The cast strips were allowed to dry under ambient conditions overnight before cutting them into square shaped films of 2.5 × 2.5 cm. The films were then stored under ambient conditions until further experiments. During preparation, ambient relative humidity was monitored using a hygrometer, and film casting was not performed if humidity exceeded 50 %, consistent with values recommended in the literature for reproducible solvent evaporation and polymer film formation.^25^

### 2.3 Physicochemical characterization of the films

Film thickness was determined with a micrometer screw gauge (Mikrometer Cocraft, Clas Ohlson, Sweden; range 0–25 mm, resolution 0.01 mm) by taking six measurements evenly distributed from the center to the edges of each film (N=3, n = 6). The morphology and structural properties of the films were visualized with Scanning Electron Microscopy (SEM) using a tabletop SEM (TM3030Plus Tabletop Microscope, Hitachi, Japan). The measurements were done at 15 kV, and the images were analyzed using ImageJ software (version 1.54 f).

A commercial spectral domain optical coherence tomography (SD-OCT) system (Telesto II, Thorlabs, Newton, NJ, USA) was employed to image the films and evaluate the optical properties of the films in static conditions and in contact with *ex vivo* porcine epithelium. The system utilized a superluminescent diode centered at 1310 nm with a spectral bandwidth of 140 nm, yielding an axial resolution of 5.5 μm in air. Imaging was carried out at a scan rate of 48 kHz using a telecentric scan lens with an effective focal length of 36 mm, a working distance of 25.1 mm, and a lateral resolution of 13 µm in air (LSM03, Thorlabs, Newton, NJ, USA). All measurements were carried out in a laboratory environment maintained at 20°C and 60 % relative humidity to ensure consistent and controlled conditions. The focal plane was adjusted to lie within the sample to optimize contrast between structural layers. All depth axes are in units of optical path length

The water content of the films was determined using a moisture analyzer (Hologen Moisture Analyzer, HE53, Mettler Toledo). Films were weighed before and after drying at 105°C until constant weight. The apparent water loss was determined as the percentage decrease in film weight after drying. Measurements were conducted in triplicate (n = 3) for both dual- and triple-layer films.

### 2.4 Cytotoxicity of films on TR146 cells

The immortalized human buccal epithelial cell line TR146 was used for experiments between passages 21 and 23. Cells were cultured in T75 polystyrene cell-culture flasks (Corning, NY, USA) under standard conditions (37_°_C, 5 % CO_2_). The culture medium consisted of DMEM/F-12 supplemented with 10 % FBS, 1 % L-glutamine, 1 % Pen/Strep, and 1 % NEAA. Culture medium was replaced three times per week. Upon reaching 80 % confluency, cells were passaged using TrypLE™ Express and seeded in new flasks in a 1:8 split ratio. Cytotoxicity was evaluated using the MTT assay, which measures cell metabolic activity based on the enzymatic reduction of the yellow tetrazolium salt (MTT) to purple formazan crystals by viable cells.^26^ TR146 cells were seeded in 96-well plates (Greiner Bio-One GmbH, Frickenhausen, Germany) at a density of 3 x 10^4^ cells/cm^2^ using the same culture medium as described above and allowed to attach for 24 h. Experiments were conducted for blank dual- and triple-layer films, with two sample groups prepared by dissolving each film (2.5 x 2.5 cm square) in 5 mL of ultrapure water. Triton X-100 (1 % v/v) served as the cytotoxic control, while untreated cells in culture medium served as the viable control. After 24 h of growth, the monolayers were washed three times with pre-warmed PBS and incubated with 150 μL of film-containing medium for 1 h at 37 _°_C in a 5 % CO_2_ incubator. Following exposure, the film solutions were removed, and cells were washed three times with pre-warmed PBS. In one experimental group, fresh culture medium was added, and the cells were allowed to recover for 24 h, while cytotoxicity was assessed immediately in the remaining group. For both groups, the media was replaced with 100 μL fresh culture medium, followed by addition of 10 μL of 12 mM MTT stock solution from the CyQuant™ MTT Cell Viability Assay Kit (Invitrogen, CA, USA), according to the manufacturer’s protocol. Plates were incubated for 4 h (37 _°_C, 5 % CO_2_). Subsequently, 100 μL of SDS-HCl solution was added to each well, and the plates were further incubated overnight (18 h) to allow for formazan solubilization. Formazan absorbance was measured spectrophotometrically at 570 nm using a microplate reader (Varioskan LUX Multimode Microplate Reader, Thermo Scientific, Waltham, MA, USA). Each condition was tested in six replicates (N=3, n=6). Results are expressed as percentage of metabolic activity relative to untreated controls, with background absorbance subtracted from all readings.

### 2.5 Osmolality measurements

Osmolality measurements were performed using an Osmomat 3000 freezing point osmometer (Gonotec GmbH, Berlin, Germany). Prior to measurements, the osmometer was calibrated with standard NaCl solution of known osmolality (300 mOsm/kg) following the manufacturer’s instructions. A 50 µL aliquot of each sample was loaded into the measurement chamber, and osmolality was determined based on the freezing point depression method. Solutions of either GLP-1-RA and GDC (10 mg/mL and 20 mg/mL, respectively) in ultrapure water as well as dissolved dual- and triple-layer films (2.5 x 2.5 cm) in 5 mL of ultrapure water, were tested. Between measurements, the chamber was cleaned and dried according to manufacturer guidelines to prevent cross-contamination. Osmolality values are reported in milliosmoles per kilogram of water (mOsm/kg H₂O).

### 2.6 *Ex vivo* evaluation of mucoadhesive properties of dual- and triple-layer films

The mucoadhesive properties of both the dual- and triple-layer films were evaluated *ex vivo* using a texture analyzer (TA.XTplus Texture Analyzer, Stable Micro Systems, Godalming, UK). The experimental parameters used for testing were adapted from literature^27^, and are summarized in Table 2. Briefly, each film was affixed to the upper probe of the instrument, which was then lowered onto the mucosal tissue and applied a predefined force of 2N. After a contact time of 60 sec allowing for adhesion, the probe was withdrawn, and the maximum detachment force (mN) required to separate the film from the tissue was recorded as a measure of mucoadhesion. From the adhesion-distance curves, the peak detachment force and the work of adhesion-corresponding to the area under the curve (AUC) were calculated.

**Table 2:** Parameters used for *ex vivo* adhesion testing of dual- and triple layered films using a texture analyzer.

| Caption | Value | Units |
| --- | --- | --- |
| Pre-test speed | 0.50 | mm/sec |
| Test speed | 0.50 | mm/sec |
| Post-Test Speed | 5.00 | mm/sec |
| Applied Force | 2.000 | N |
| Return Distance | 20.000 | mm |
| Contact time | 60 | sec |

Freshly excised epithelium from porcine buccal tissue, was carefully glued to the base of a Petri dish, taking care to maintain a smooth, flat surface while avoiding any stretching or mechanical damage to the tissue. The sample was kept hydrated by applying 200 µL of a 3 % (w/v) mucin solution in PBS. This ensured continuous moisture throughout the experiment, simulating physiological conditions.^28^ All measurements were performed under consistent environmental conditions. A total of 15 replicates for each type of film were analyzed to ensure statistical reliability (N=15).

### 2.7 UPLC analysis

Chromatographic analysis of GLP-1-RA was performed on a Waters ACQUITY UPLC H-Class system (Waters Corporation, Milford, MA, USA), equipped with a photodiode array (PDA) detector. Separation was achieved using an ACQUITY UPLC BEH C18 column (50 × 2.1 mm, 1.7 µm particle size) maintained at 30 °C. The mobile phase consisted of eluent A (0.1% trifluoroacetic acid in water) and eluent B (0.09% trifluoroacetic acid in acetonitrile). The flow rate was set to 0.7 mL min⁻¹, and the injection volume was 2 µL for the release study or 20 µL for the permeability study. The column effluent was monitored at a wavelength of 214 nm. Samples were maintained at 7 °C in the autosampler prior to injection. The gradient elution program is summarized in Table 3. The total run time was 5.1 min, and the analyte of interest (GLP-1-RA) eluted at approximately 3.2 min under these conditions.

**Table 3:** Gradient elution program for UPLC analysis.

| Time | Flow | %A | %B |
| --- | --- | --- | --- |
| Initial | 0.7 | 70 | 30 |
| 3.2 | 0.7 | 45 | 55 |
| 3.6 | 0.7 | 10 | 90 |
| 3.7 | 0.7 | 5 | 95 |
| 4.0 | 0.7 | 5 | 95 |
| 4.1 | 0.7 | 70 | 30 |
| 5.1 | 0.7 | 70 | 30 |

### 2.8 *In vitro* GLP-1-RA release from dual- and triple-layer films

*In vitro* release of GLP-1-RA and GDC from the films (squares of 2.5 x 2.5 cm) was evaluated using 6-well plates. Films were placed at the bottom of each well with ML facing upward for triple-layer films and AL facing upward for dual-layer films, ensuring direct contact with the release medium. PBS (pH 7.4; 5 mL) was used as the release medium in all experiments. The release studies were conducted at 37 °C, without agitation to assess passive diffusion behavior.

At predetermined time intervals, aliquots of the release medium (200 μL) were withdrawn for analysis and replaced with fresh PBS. The samples collected were analyzed for drug content using ultra-performance liquid chromatography (UPLC), and the cumulative release of GDC and GLP-1-RA over time was calculated. Each experiment was performed in quadruplicate (N = 4).

### 2.9 Circular dichroism analysis of peptide stability post-casting

Circular dichroism (CD) spectroscopy was employed to evaluate the structural integrity of the peptide following the film casting process. Films (squares of 2.5 x 2.5 cm) containing the peptide were dissolved in ultrapure water (5 mL) to release GLP-1-RA into solution. As a control, blank films (without GLP-1-RA and GDC) were dissolved at identical conditions to account for any background signal from the film matrix. CD measurements were performed using a spectropolarimeter (Applied Photophysics Chirascan, Leatherhead, United Kingdom) at room temperature. Spectra (n=5) were recorded over a wavelength range of 180-250 nm using a quartz cuvette with a path length of 1 mm. The resulting spectra of peptide-containing samples were corrected by subtracting the signal from the blank film solution, followed by baseline correction. Data points with absorbance values greater than 2 were excluded from analysis to ensure accuracy and reliability of the results.

### 2.10 *Ex vivo* penetration and post-exposure tissue analysis

*Ex vivo* permeability studies were conducted using porcine buccal tissue obtained from healthy experimental pigs (Danish Landrace/Yorkshire/Duroc, approximately 30-40 kg) provided by the Department of Veterinary and Animal Sciences, University of Copenhagen. Buccal cheek tissue was collected immediately post-euthanization and stored in PBS on ice until use (utilized on the same day as harvesting). The tissue was trimmed to remove underlying connective layers. The cheeks were then placed in a water bath of 70°C for 4 min followed by peeling the epithelium off. The epithelium was then mounted in custom-fabricated permeable support systems, with the film positioned on the surface. Round films of 10 mm diameter were used for the study. For triple-layer films, ML was placed in contact with the tissue, while for dual-layer films, AL faced the tissue to ensure unidirectional drug release toward the epithelium. The receptor chamber was filled with 1.5 mL of PBS, maintained at 37 °C throughout the experiment. Penetration studies were conducted over a 3-h period with 200 μL samples collected from the receptor compartment at predetermined time points (15, 30, 60, 120 and 180 min). Following each sampling, an equal volume of fresh pre-warmed PBS was added. Collected samples were diluted with PBS and analyzed by UPLC to quantify the penetration of GDC and GLP-1-RA. Each condition was tested in quadruplicate (N = 4). As a control, a parallel experiment was conducted using a Hilltop chamber (Hilltop chambers, 19 mm, Cliantha Research, Saint Petersburg, FL, USA). Hilltop chambers are small diffusion cells designed to hold liquid formulations on the tissue surface in a controlled manner.^29^ They enable direct contact between the tissue and the solution while preventing sample leakage or spreading. Here, equivalent amounts of GDC and GLP-1-RA (matching the content of the films) were placed directly in the donor compartment without a film matrix (N=4). The solution was pipetted on the cotton which was allowed to soak prior to placement on the tissue. The cumulative amount of GDC and GLP-1-RA penetrating the epithelium over time was calculated.

For monitoring epithelial barrier integrity during the experiments, fluorescein isothiocyanate dextran (FD70, 100 μg/mL) was added to the donor compartment. Additionally, transepithelial electrical resistance (TEER) was measured before and after the experiment to assess changes in tissue barrier properties. The difference in TEER values between 0 and 180 min was calculated and expressed as a percentage to assess changes in epithelial barrier integrity under different experimental conditions.

Following the permeability experiments, selected tissue samples were immediately placed in 10 % v/v neutral buffered formalin at a ratio of approximately 10:1 (fixative volume: tissue volume) and fixed at room temperature for 24 h. Tissue samples were fixed in 10% NBF for at least 24 hours. Following fixation, tissues were processed, dehydrated and infiltrated with paraffin, using the Excelsior AS Tissue Processor and embedded in paraffin blocks on the HistoCore H. Paraffin sections were cut at 3 μm using the HistoCore AUTOCUT microtome. Slides were stained with H&E using Meyer’s Hematoxylin and Eosin Y 1% alcoholic solution, diluted to 0.2% in 96% ethanol with 2.5 mL glacial acetic acid. Finally, slides were mounted with Pertex for microscopic analysis.^30^

Other samples were washed in 4 % w/v paraformaldehyde (PFA) in PBS and imaged using coherent anti-Stokes Raman scattering (CARS) microscopy using a Leica TCS SP8 CARS microscope equipped with a set of CARS 2000 filters, HC PL IRAPO 40×/1.10 WATER objective (Leica, Germany). CARS was performed with a fixed Stokes laser beam of 1032 nm while the wavelength of pump laser was tuned to 797.1 nm. This generates a CARS signal corresponding to 2850 cm^-1^ Raman shift from -CH- bond in lipid to assess lipid content in the cell membranes, allowing for evaluation of any film-induced alterations to tissue morphology.

### 2.11 Data and statistical analysis

The data analysis was conducted using GraphPad Prism (Version 9.4.1 (681), Insight Partners, GraphPad Holdings, LLC, New York City, NY, USA). Results are presented as mean ± standard deviation (SD), where N denotes the number of independent film samples (physical replicates), while n represents the number of repeated measurements performed on each sample. Statistical significance was assessed with GraphPad Prism, employing a one-way analysis of variance (ANOVA) for comparisons among three or more independent groups, with a significance threshold set at *p < 0.05, **p < 0.01, and ***p < 0.001. Schematics were created using BioRender.com. Image analysis and processing were performed with ImageJ software (version 1.53t, National Institutes of Health, USA).

## 3. RESULTS & DISCUSSION

### 3.1 Film characterization

Dual- and triple-layer films were successfully fabricated using slot-die coating (Fig. 1A and B). Materials for each layer were selected based on their specific functional roles. The ML (present only in the triple-layer films) promotes adhesion to the buccal tissue. In the dual-layer films, the AL functions as both a mucoadhesive and a drug-loaded layer, whereas in the triple-layer films, it serves solely as a drug-loaded layer. The BL functions as a barrier to enable unidirectional release and controlled film disintegration, enhancing patient comfort. The composition and functional roles of all components are summarized in Table 4.

**Table 4:**
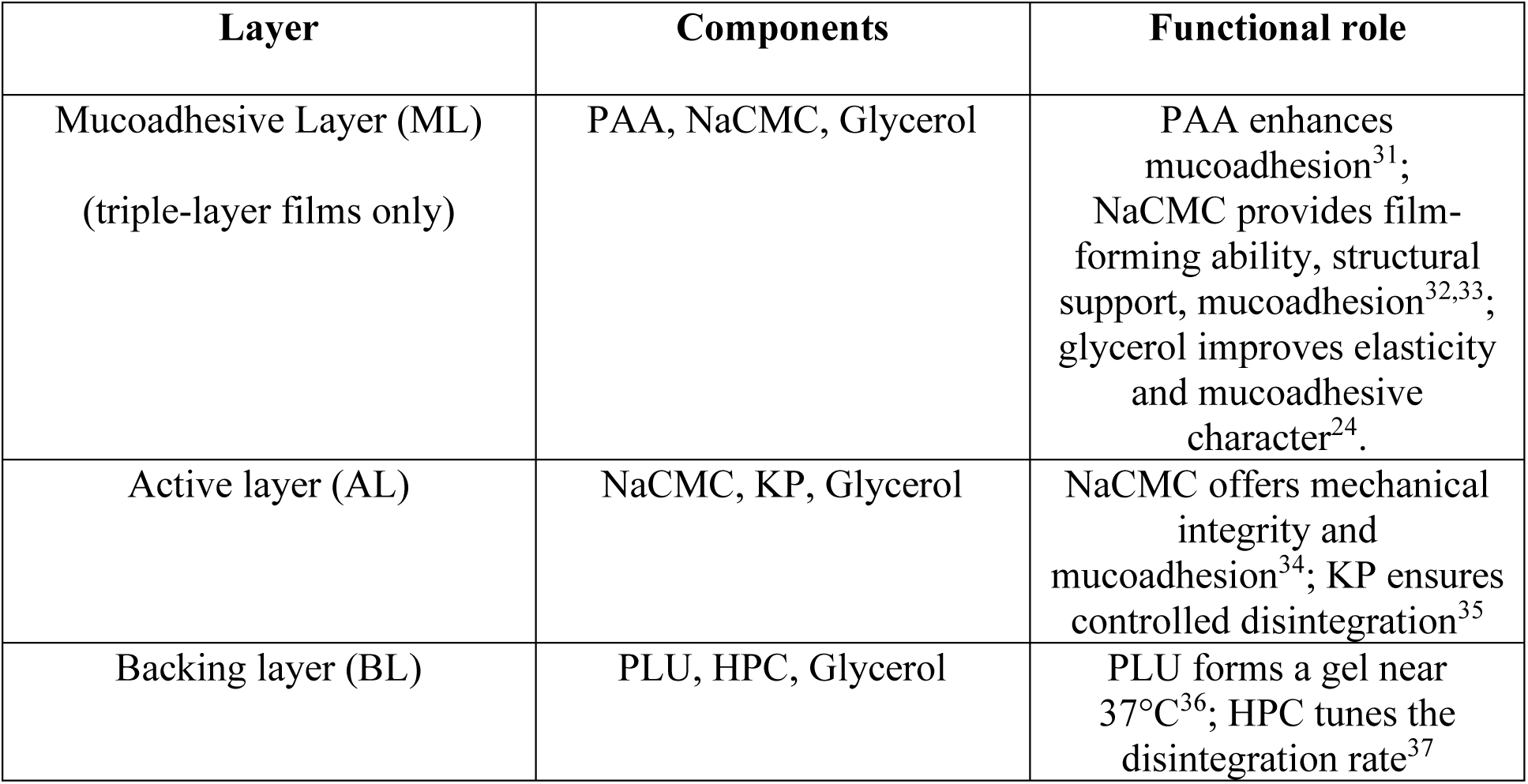
Summary of the materials used in each layer of the multilayer buccal films and their corresponding functional roles, including mucoadhesion, drug loading, controlled release, and film dissolution.

Slot-die coating has previously been used to fabricate single- and dual-layer films.^24,38^ Investigating the effect of casting order showed that when the BL was cast first, film formation was suboptimal, likely due to its composition. Conversely, casting the ML and AL first allowed the BL to spread homogeneously on top, yielding well-formed multilayer films as schematically shown in Fig. 1A.

The films were characterized by SEM, and their overall thickness was measured using a micrometer. The average total thicknesses were 364.0 ± 17.0 µm for the dual-layer films and 465.3 ± 24.6 µm for the triple-layer films (N=3, n=10). Using OCT, distinct layers were observed in both dual- and triple-layer films (Fig.1B, C). In addition, there were clear differences in textures for each layer evident from the SEM images (Fig.1B, C). Layer-specific thicknesses were further analyzed using SEM imaging. The dual-layer films had a total thickness of 339.40 ± 10.24 µm, consisting of an AL measuring 151.28 ± 5.66 µm and a BL measuring 192.60 ± 4.59 µm. The triple-layer films were thicker, with a total thickness of 487.0 ± 36.5 µm, comprising a ML of 171.7 ± 15.7 µm, an AL of 207.7 ± 16.3 µm, and a BL of 107.6 ± 4.5 µm. In the literature, buccal films typically range from 50 to 1,000 µm in thickness.^39,40^ The dual- and triple-layer films developed in this study measured approximately 339 µm and 487 µm, respectively, placing them well within the typical range for buccal formulations. For comparison, commercial orodispersible tablets often have thicknesses of 2-5 mm and are generally well tolerated.^41–43^ Therefore, the present multilayer films should not inherently compromise patient compliance or usability, particularly given their flexible and mucoadhesive nature.

The moisture content of the multilayer films was determined to evaluate their water content. The dual-layer films exhibited a moisture content of 3.85 ± 0.91 %, whereas the triple-layer films showed a higher moisture content of 6.37 ± 0.09 % (n=3). The increase in moisture content with additional layers may influence the films’ hydration and swelling behavior, potentially affecting the release properties of GLP-1-RA and GDC. Higher moisture in the triple-layer films could enhance film flexibility and promote faster hydration of the individual layers, which may facilitate more efficient sequential release of GDC and GLP-1. However, elevated moisture levels may also increase the risk of GLP-1 degradation, potentially compromising the stability and efficacy of the active pharmaceutical ingredients.^44–46^

For evaluation of the cytocompatibility of the polymeric films, experiments were performed using TR146 cells. These cells are a human buccal epithelial cell line commonly employed as an *in vitro* model of the buccal epithelium due to their ability to form stratified epithelial layers and mimic the barrier properties of the buccal epithelium.^47^ Solutions of dissolved blank dual-and triple-layer films were incubated with TR146 cells for 1 h and 24 h. Cell viability assays showed that exposure to either of the film types did not reduce cell viability at either time point compared to the control (Fig. 2), demonstrating that the polymeric matrices themselves are non-toxic and suitable for oral mucosal applications. This observation aligns with existing literature, which reports that polymeric matrices designed for oral mucosal applications are typically non-toxic and biocompatible.^48^

**Figure 2:**
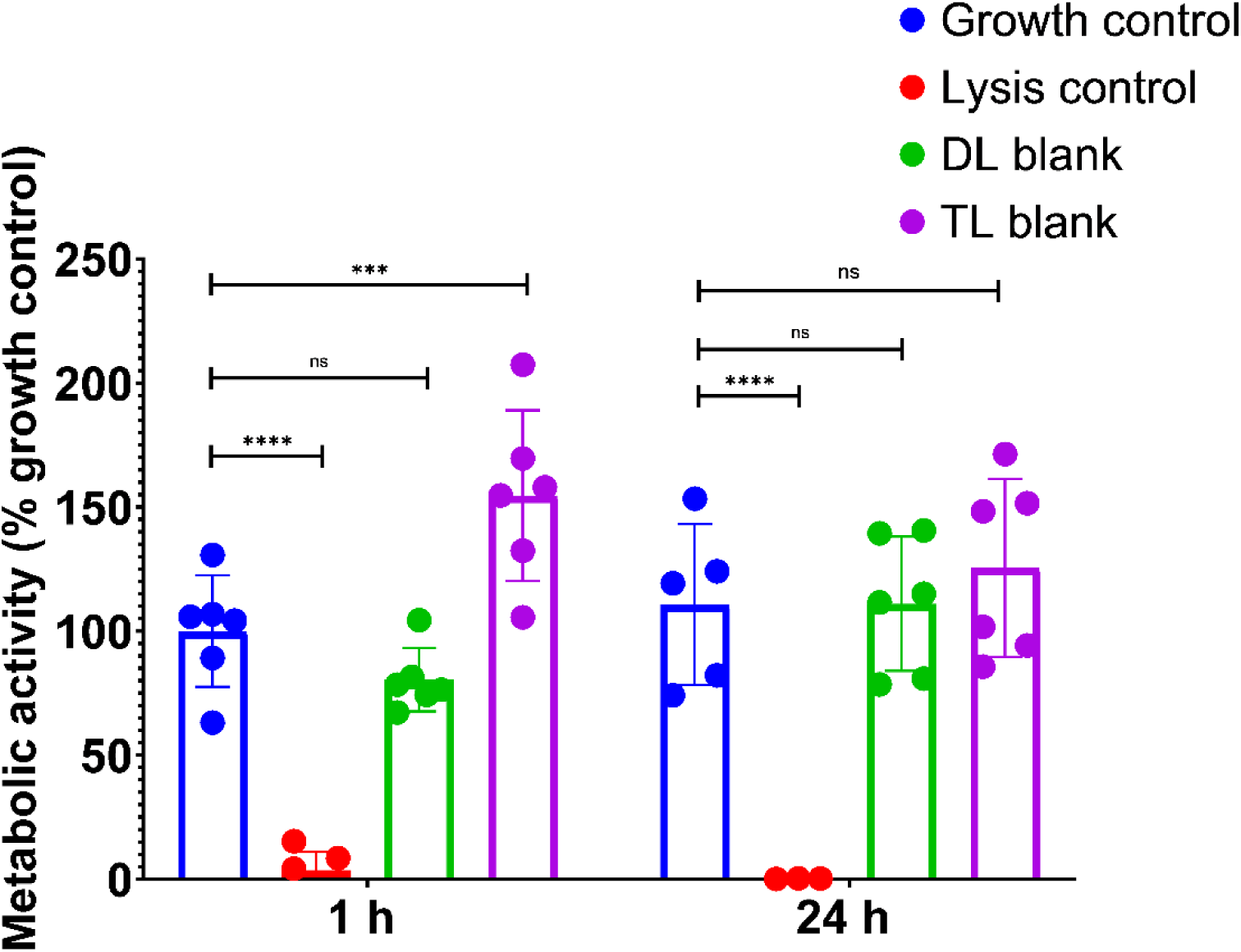
Cytotoxicity assessment via MTT test of blank films demonstrated that the polymeric matrices themselves were non-toxic to TR146 cells (DL: dual layer, TL: triple layer). Data represents the mean ± SD with n=6. Statistical significance was determined using one-way ANOVA followed by Dunnett’s multiple comparisons test. p < 0.05 was considered statistically significant; asterisks indicate significant differences compared to the control group.

### 3.2 Mucoadhesive properties of dual- and triple-layer films

The mucoadhesive properties of the films were quantitatively evaluated *ex vivo* using a texture analyzer at physiologically relevant and controlled conditions (Fig. 3A). Round films of 10 mm diameter were used. Representative force-distance curves for both film types are presented in Fig. 3B, highlighting the detachment profile. As shown in Fig. 3C, the average maximum detachment force was 1571.3 ± 126.6 mN for the dual-layer films and 1562.9 ± 388.2 mN for the triple-layer films. The corresponding work of adhesion values were 1.96 ± 0.23 N·mm and 2.02 ± 0.17 N·mm, respectively. No statistically significant differences were observed between the dual- and triple-layer films, indicating comparable mucoadhesive performance across both delivery systems. The adhesive strengths reported in the literature for mucoadhesive films are typically between 0.4 and 1.1 N/cm² depending on polymer type and composition (e.g., PVP, chitosan, PVA blends).^49–53^ However, direct comparison is not possible due to significant variations in experimental protocols, such as differences in contact force, deformation or withdrawal rate, contact time, and film area. Consequently, while present data suggest superior mucoadhesive performance, definitive comparison with the literature remains constrained by methodological heterogeneity.

**Figure 3:**
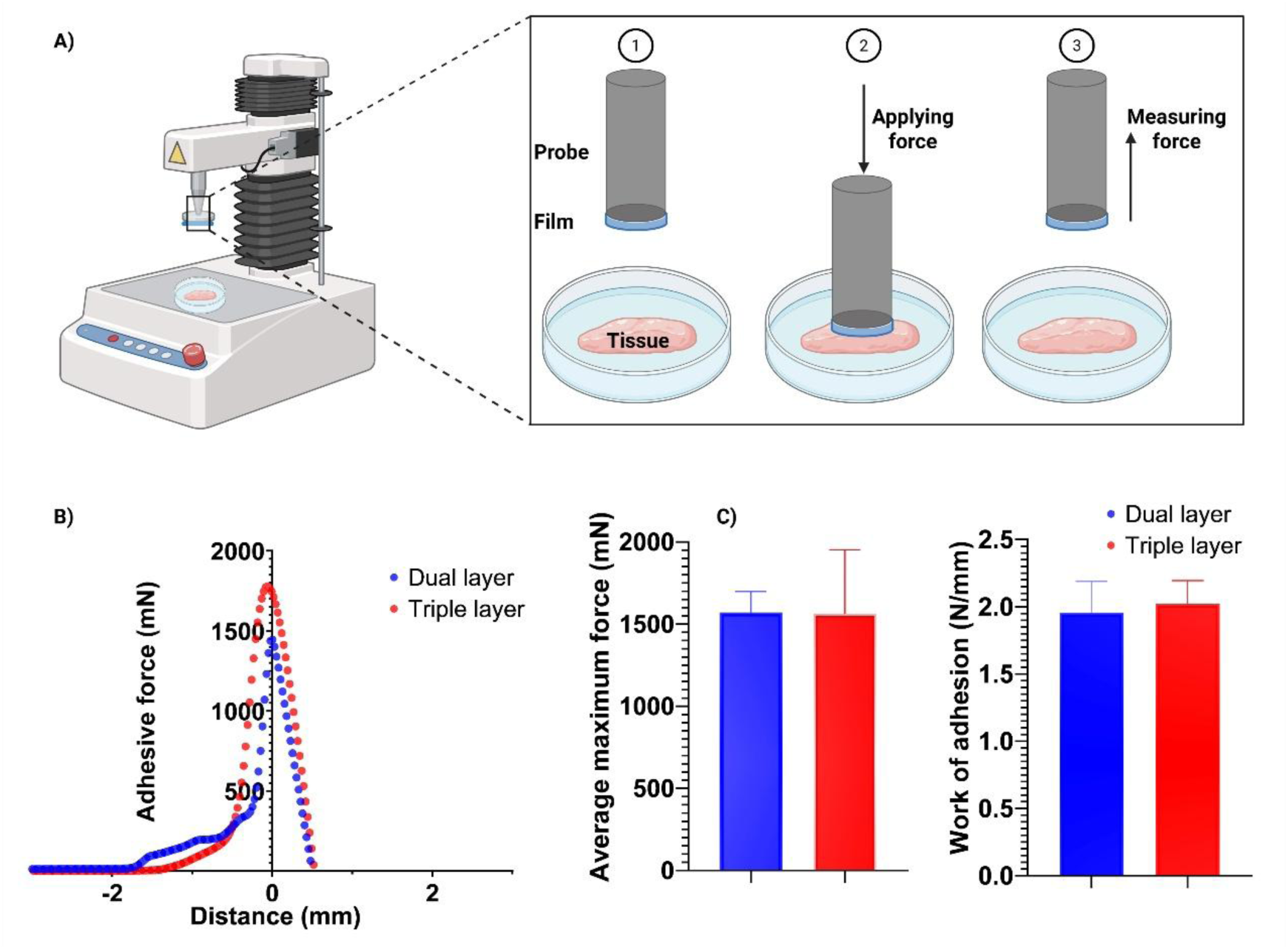
Evaluation of mucoadhesive properties of the films. A) Schematic of the experimental setup using a texture analyzer for detachment force measurements. B) Representative force–distance curves for dual-layer and triple-layer films. C) Quantitative comparison of average maximum detachment force and work of adhesion for both film types. Data is presented as mean ± SD, N=15.

OCT imaging was used to monitor the hydration and swelling behavior of the films in contact with *ex vivo* porcine tissue (Fig. 4). In the dual-layer films, the AL in direct contact with the tissue exhibited rapid swelling, driven by water uptake at the tissue interface. In the triple-layer films, swelling occurred predominantly in the ML, which is the one in contact with the tissue, while the AL remained largely unchanged. In both film types, the layer directly contacting the tissue shows the greatest swelling, regardless of its composition, and the ML continues to swell over time. In contrast, swelling of the AL depends on its position within the multilayer structure. Overall, the films exhibit similar swelling patterns, which aligns with their comparable mucoadhesion behavior observed. In both film types, the BL swelled gradually, consistent with its design for controlled hydration and disintegration.

**Figure 4:**
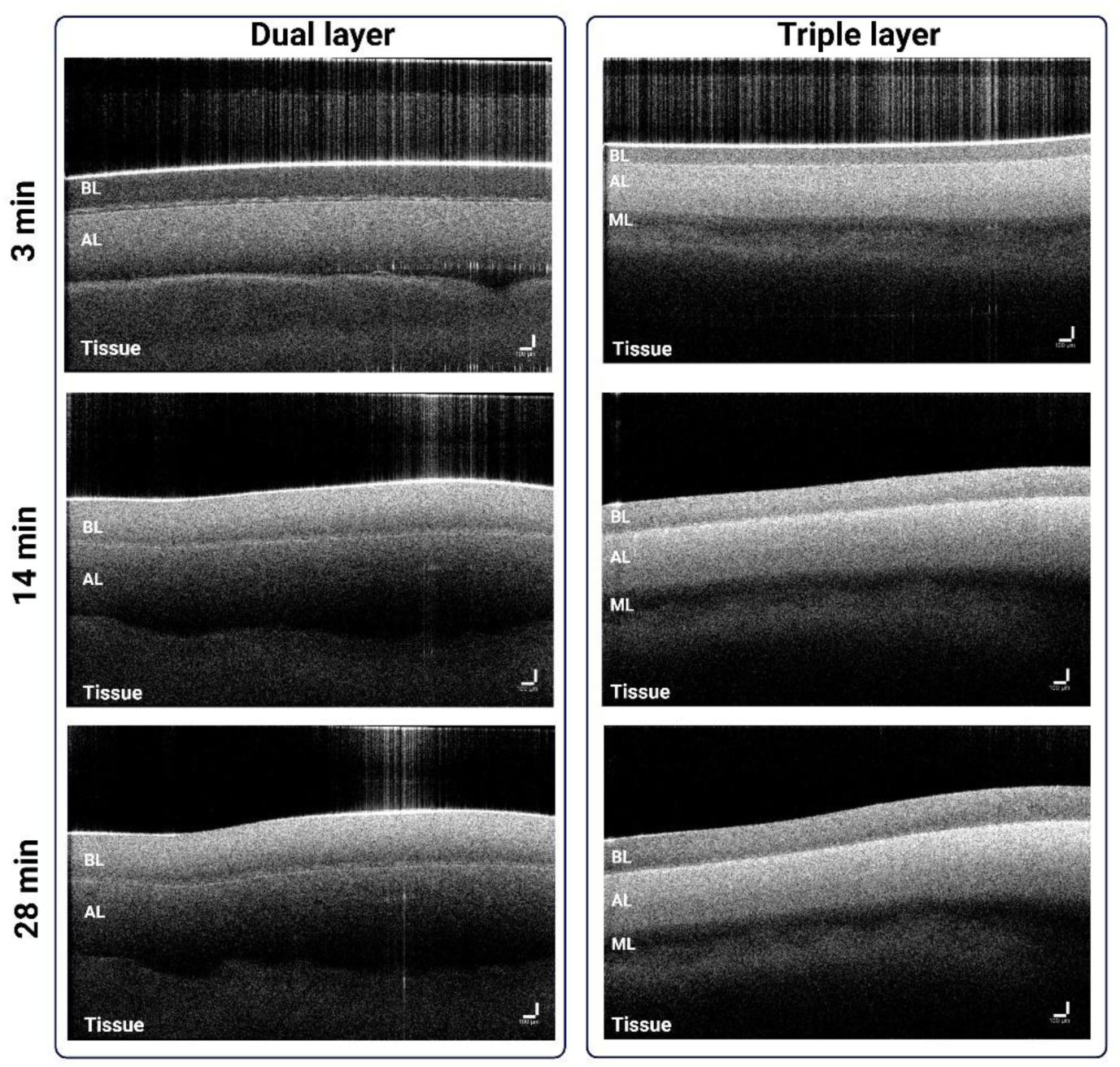
Optical coherence tomography (OCT) representative images of dual- and triple-layer films in contact with *ex vivo* porcine tissue for 3-, 14- and 28-min. Experiments performed at controlled temperature environment (20°C). (ML: mucoadhesive layer, AL: active layer, BL: backing layer). Scale bars represent 100 μm.

### 3.3 *In vitro* drug release and GLP-1-RA stability evaluation

To assess the structural integrity of the GLP-1-RA following the casting process, CD spectroscopy was performed on film samples dissolved in water. Spectra were recorded for the individual components (GDC and GLP-1-RA), as well as their mixture. To confirm that the film matrix did not interfere with the measurements, blank films were cast, dissolved under identical conditions, and their spectra recorded. The resulting overlaid spectra are presented in Fig. 5A, with the respective absorbance plotted in Fig. 5B. The CD spectrum of the GLP-1-RA in solution exhibits characteristic features of an α-helical conformation, with a strong positive peak near 190 nm and two distinct negative minima at approximately 208 nm and 222 nm.^54,55^

**Figure 5:**
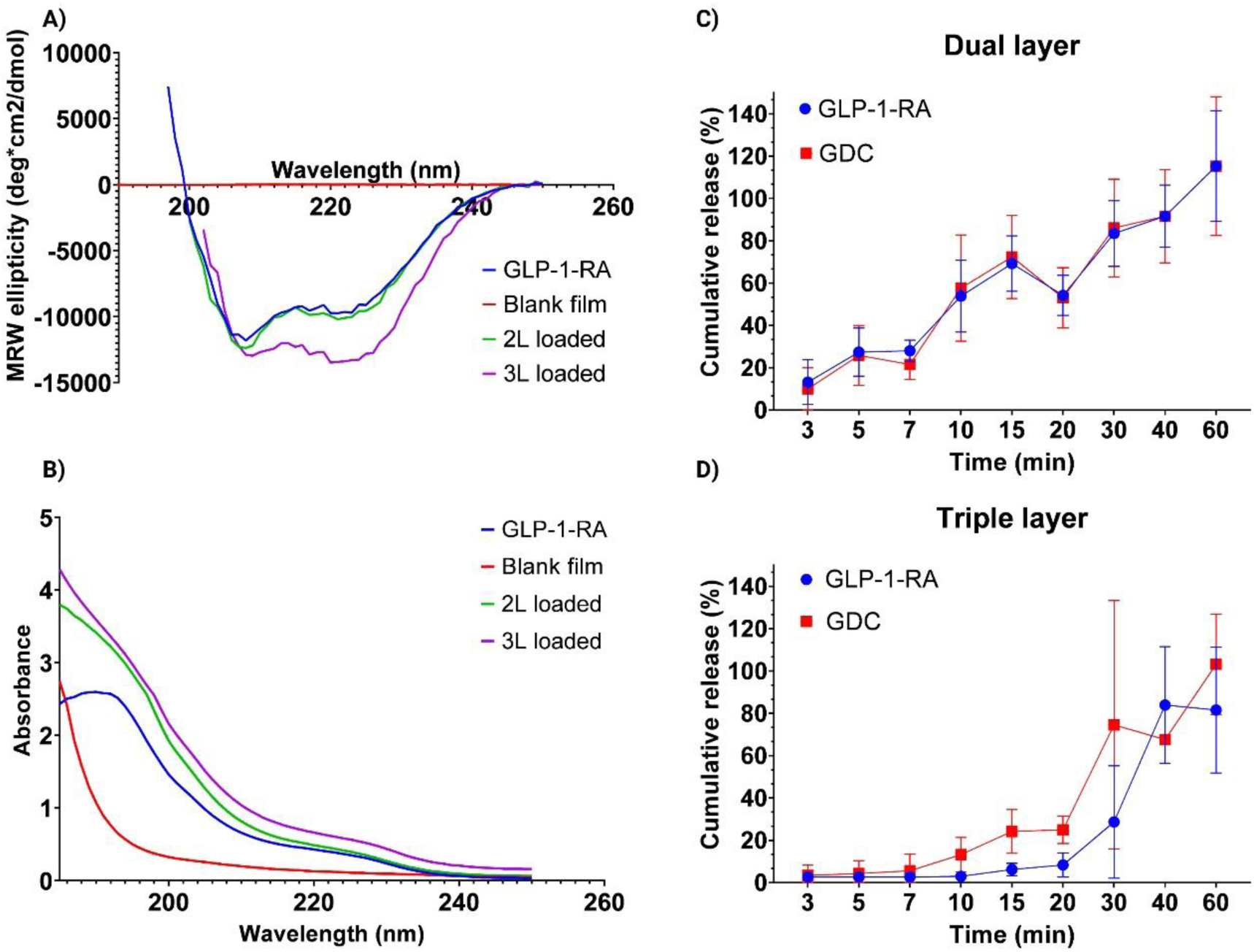
Evaluation of peptide integrity and drug release from dual and triple layer films. (A) Circular dichroism (CD) spectra of GDC, GLP-1-RA, their mixtures, and dissolved blank films, indicating preservation of secondary structure after film casting and the absence of spectral contributions from the film matrix (n=5). (B) Corresponding absorbance spectra, presented to ensure that measurements were acquired within the reliable CD detection range (absorbance <2). (C,D) Cumulative release profiles of GDC and GLP-1-RA from dual- and triple-layer films under static conditions (PBS, 37°C), respectively. Data represents mean ± SD, N=4.

To assess the structural integrity of the GLP-1-RA after undergoing the casting process for the dual- and triple-layer films, CD measurements were repeated. Data below 200 nm are not plotted, as the absorbance exceeded 2 for both films. The overall spectral shape of the dual-layer film was largely preserved, suggesting that the peptide retained its predominantly α-helical structure post-casting. In contrast, the triple-layer sample showed noticeable differences, including higher overall absorbance in the 220–250 nm range, which may indicate contributions from other absorbing compounds. These observations suggest that the altered CD profile of the triple-layer sample may not be solely due to structural changes in the GLP-1-RA.^56,57^

The *in vitro* release profiles of GDC and GLP-1-RA from both dual- and triple-layer films (squared 2.5 x 2.5 cm) are shown in Fig. 5C and 5D, respectively. During solution preparation, equal amounts of GLP-1-RA and GDC were incorporated in both film types. Quantification showed that 12.80 ± 1.98 mg/film and 11.85 ± 1.98 mg/film of free GLP-1 RA were detected in the dual- and triple-layer films, respectively. The free amount of GDC that was measured was 8.61 ± 0.29 mg/film in the dual-layer films and 18.46 ± 7.15 mg/film in the triple-layer films. During static conditions, dual-layer films showed a rapid release, with GLP-1-RA reaching 13.2 % ± 10.6 % after 3 min and 115.3 % ± 26.1 % after 60 min. GDC followed a similar trend. In contrast, triple-layer films exhibited a slower initial release, with GLP-1-RA at 2.6 % ± 0.06 % at 3 min, gradually increasing to 81.6 % ± 29.8 % by 60 min; GDC showed comparable sustained release. Notably, triple-layer films had a delayed but distinct increase between 20- and 40-min. Release kinetics of GLP-1-RA and GDC from dual- and triple-layer films were analyzed using common mathematical models. For the dual-layer films, both compounds showed the best fit to the Korsmeyer-Peppas model, indicating that drug release followed an anomalous transport mechanism, governed by a combination of diffusion and matrix dissolution.^58^ In the case of the triple-layer, the system exhibited different release mechanisms for each compound: GLP-1-RA release followed first-order kinetics, indicating a release rate dependent on the remaining drug concentration, while GDC release had the best fit to the Korsmeyer-Peppas model, implying a combination of diffusion and other processes such as matrix relaxation or erosion.^58^ Overall, these results demonstrate the functional impact of film architecture. In the dual-layer films, the release curves of GLP-1-RA and GDC were nearly identical across all time points, indicating that both compounds diffused out simultaneously. In contrast, the triple-layer films exhibited a clear sequential pattern; with GDC released first in a pronounced early phase, followed by a later increase in GLP-1-RA, confirming that the multilayer design effectively separates and modulates the release of the two compounds. This temporal separation highlights that the films’ architecture enables programmable release that is not achievable with the simpler configurations. These release differences align with the OCT swelling behavior. In the dual-layer films, rapid swelling of the AL at the tissue interface created an immediate hydration front, explaining the fast and simultaneous release of both compounds. In contrast, the triple-layer films showed swelling concentrated in the ML while the AL remained largely unchanged early on, producing a slower, more structured hydration pattern. This corresponds directly to the staggered release profile of the triple-layer system, with GDC released first and GLP-1-RA following later, demonstrating how the multilayer architecture governs diffusion and timing of release. Previous studies have shown that the release profile from buccal films can vary significantly depending on formulation characteristics such as polymer composition, drug loading, and film thickness. Rapid release has often been associated with more soluble drug forms and hydrophilic matrices, while sustained release profiles are commonly linked to more structured systems.^52,59^ As shown by previous studies, static (no stirring) release conditions may better mimic the low-fluid environment of the oral cavity, particularly relevant for buccal applications.^52,60,61^

### 3.4 Impact of film formulation on buccal penetration and epithelial morphology

The *ex vivo* permeability of GLP-1-RA and GDC was evaluated using fresh porcine buccal epithelium. Both compounds exhibited time-dependent penetration, with faster and higher transport from the triple-layer film compared to the dual-layer (Fig. 6A). For GLP-1-RA, early penetration was minimal (<0.1 % up to 30 min), gradually rising after 60 min to reach 0.15 % (dual-layer) and 0.37 % (triple-layer) at 180 min. In contrast, GDC permeated more rapidly and extensively with levels of 3.8 % (dual layer) and 8.3 % (triple layer) at 15 min, increasing to 10.6 % and 25.1 % at 180 min, respectively. These data highlight two key findings: (i) the triple layer system consistently permitted higher flux for both compounds, and (ii) GDC penetrated far more readily than GLP-1-RA. A plausible explanation for the enhanced flux in the triple layer system is the spatial separation of the compounds within the film. The mucosa is likely exposed first to the layer containing GDC, allowing it to act on the tissue before GLP-1-RA release from the subsequent layer. This sequential exposure may prime the tissue, enhancing penetration.^62^ In the dual layer system, both GDC and GLP-1-RA are released simultaneously, so the enhancer has limited opportunity to act before GLP-1-RA diffusion begins. This difference in release order likely contributes to the higher penetration observed with the triple layer films. Additionally, the unidirectional release of GDC in the triple-layer film may lead to a higher local concentration at the tissue interface, whereas in the dual-layer system some GDC could diffuse away from the opposite side, reducing its local effect.

**Figure 6:**
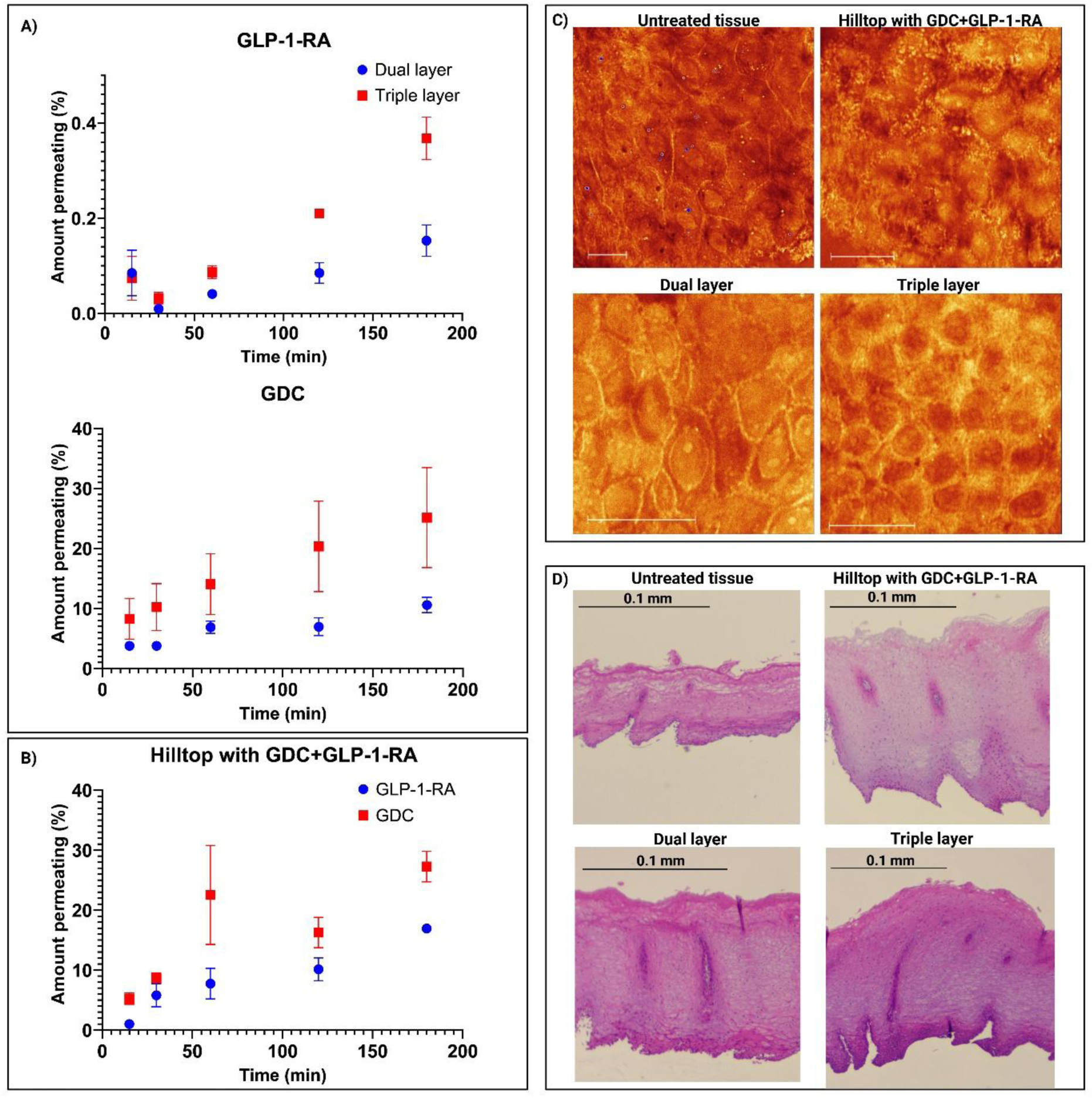
*Ex vivo* evaluation of GLP-1-RA and GDC delivery from dual- and triple-layer films. A) Time-dependent cumulative penetration of GLP-1-RA and GDC across porcine buccal mucosa from dual- and triple-layer films. B) Cumulative penetration from the Hilltop chamber control, where compounds were applied without polymeric incorporation. C) Representative CARS microscopy images of epithelial tissue after exposure to each formulation, showing differences in cell morphology and integrity. Scale bars represent 20 μm. D) H&E-stained porcine buccal epithelium corresponding to the treatments in (C), confirming structural changes in the epithelium associated with formulation type and osmolality. Scale bars represent 0.1 mm. Data is presented as mean ± SD, N=4.

In this study, Hilltop chambers were used to allow exposure of the tissue to the formulation components without a polymeric matrix, while keeping the solution securely in place, as shown in (Fig. 6B). In this case, where GLP-1-RA and GDC were applied directly without incorporation into a polymer matrix, both compounds permeated more rapidly and extensively; GLP-1-RA reached 16.9 ± 0.65 % and GDC 27.3 ± 5.11% at 180 min. This suggests that polymeric films reduce the freely available fraction of the compounds, likely due to binding interactions, limiting effective penetration.^63^ Thus, while film-based delivery provides more controlled release, polymer–compound interactions modulate flux, emphasizing the importance of film design in achieving optimal delivery profiles.

TEER measurements over 180 min showed minimal changes in untreated tissue (10%) and slightly higher changes in tissue exposed to the blank Hilltop chamber (16%), indicating that the experimental setup itself can modestly influence tissue integrity even in the absence of any formulation. Inclusion of GDC with GLP-1-RA caused a pronounced TEER drop (58.6 %), consistent with GDC’s role as a penetration enhancer that transiently increases membrane fluidity.^64^ Dual- and triple-layer films reduced this effect (55.7 % and 48.4 %, respectively), indicating that the polymer layers help preserve barrier integrity, consistent with the cytocompatibility observed in cell studies (Fig. 2). No FD70 was detected in the receptor compartment, confirming that paracellular leakage of FD70 did not occur.

To better understand the effects of the films on tissue morphology, CARS microscopy and histological analyses were performed following the experiment (Fig. 6C, D). Untreated tissue exhibited tightly packed, well-structured epithelial cells.^65^ Exposure to GDC and GLP-1-RA using the Hilltop chambers showed the highest penetration, but caused extensive cellular disruption and loss of structure, consistent with its extremely hypotonic osmolality (10 mosmol/kg) and rapid osmotic swelling. Dual-layer films (146 mosmol/kg) induced moderate swelling and deformation, correlating with lower penetration. By comparison, triple-layer films (380 mosmol/kg) maintained more compact cell morphology while achieving higher GLP-1-RA flux, indicating that sequential layering enabled the enhancer to prime the tissue efficiently while minimizing cellular stress. These observations demonstrated that film design and osmolality influence penetration and epithelial morphology. Compared with isotonic oral mucosa models (∼300 mosmol/kg), the extremely hypotonic GDC and GLP-1-RA solution likely overwhelms osmotic mechanisms, causing rapid swelling, structural disruption, and markedly increased penetration.^66,67^ In contrast, the hypertonic triple-layer film preserves epithelial structure while still promoting enhanced flux. Histology supported the CARS findings: untreated tissue showed normal architecture^65^, the Hilltop control caused extensive disruption^68^, while dual- and triple-layer films maintained normal tissue structure.^69^

Overall, these results confirm that film formulation and osmolality together modulate both penetration and epithelial health, highlighting that sequential layering can enhance delivery while minimizing tissue damage, which is an important consideration for safe oromucosal drug administration.^70^

## CONCLUSIONS

This study established multilayer polymeric films, produced by a slot-die coater, as a highly promising strategy for oromucosal peptide delivery, offering unique control of release kinetics and epithelial penetration. Dual-layer films enabled rapid GLP-1-RA release, whereas triple-layer architectures achieved sustained, sequential delivery, effectively enhancing the buccal permeability of both GLP-1-RA and the penetration enhancer GDC. The films combined robust structural integrity, excellent epithelial compatibility, and good mucoadhesion, while preserving GLP-1-RA’s α-helical conformation post-fabrication. Notably, both films contain physiologically relevant doses of GLP-1-RA and GDC, comparable to marketed products.^71^ These findings highlight, for the first time, the potential of tailored multilayer film architecture to overcome the limitations of conventional monolayer systems, positioning them as a patient friendly, scalable and clinically relevant platform for peptide therapeutics.

## ACKNOWLEDGMENTS

We would like to thank Dr. Sahil Malhotra of University College Dublin for his work on establishing the *ex vivo* porcine tissue dissection protocol as well as Mr. Sandeep Karki, Dr. Muhammad Ijaz and Professor David Brayden of University College Dublin for establishing and sharing their work on the 3D printed custom made permeable support devices, as well as providing the TR146 cells.

## FUNDING

This work was supported by the European Union’s Horizon Europe research and innovation program, 101091765 - BUCCAL-PEP - HORIZON-CL4−2022-RESILIENCE-01. Views and opinions expressed are however those of the author(s) only and do not necessarily reflect those of the European Union. Neither the European Union nor the granting authority can be held responsible for them.

## Notes

### Competing Interest Statement

The authors have declared no competing interest.

